# The Validity of the Coalescent Approximation for Large Samples

**DOI:** 10.1101/170928

**Authors:** Andrew Melfi, Divakar Viswanath

## Abstract

The Kingman coalescent, widely used in genetics, is known to be a good approximation when the sample size is small relative to the population size. In this article, we investigate how large the sample size can get without violating the coalescent approximation. If the haploid population size is 2*N*, we prove that for samples of size *N*^1/3−*ϵ*^, *ϵ* > 0, coalescence under the Wright-Fisher (WF) model converges in probability to the Kingman coalescent in the limit of large *N*. For samples of size *N*^2/5−*ϵ*^ or smaller, the WF coalescent converges to a mixture of the Kingman coalescent and what we call the mod-2 coalescent. For samples of size *N*^1/2^ or larger, triple collisions in the WF genealogy of the sample become important. The sample size for which the probability of conformance with the Kingman coalescent is 95% is found to be 1.47 × *N*^0.31^ for *N* ∈ [10^3^, 10^5^], showing the pertinence of the asymptotic theory. The probability of no triple collisions is found to be 95% for sample sizes equal to 0.92 × *N*^0.49^, which too is in accord with the asymptotic theory.

Varying population sizes are handled using algorithms that calculate the probability of WF coalescence agreeing with the Kingman model or taking place without triple collisions. For a sample of size 100, the probabilities of coalescence according to the Kingman model are 2%, 0%, 1%, and 0% in four models of human population with constant *N*, constant *N* except for two bottlenecks, recent exponential growth, and increasing recent exponential growth, respectively. For the same four demographic models and the same sample size, the probabilities of coalescence with no triple collision are 92%, 73%, 88%, and 87%, respectively. Visualizations of the algorithm show that even distant bottlenecks can impede agreement between the coalescent and the WF model.

Finally, we prove that the WF sample frequency spectrum for samples of size *N*^1/3−*ϵ*^ or smaller converges to the classical answer for the coalescent.

## Introduction

The Kingman coalescent (Kingman, 1982a,b) is a mathematical model of the genealogy of *n* haploid samples. If *k* lineages are present in some earlier generation, those lineages induce a partition of the *n* current samples into *k.* For convenience, we will refer to lineages present in earlier generations as ancestral samples.^1^

Kingman’s motivation in deriving the coalescent (Kingman, 1982a,b) was to gain an understanding of the structure of Ewens’ sampling formula (Ewens, 1972, Durrett, 2008). The coalescent gives an almost instantaneous derivation of Ewen’s sampling formula, and Ewens’ sampling formula is exact under the coalescent approximation. The coalescent is perfectly memoryless because at every coalescence exactly two ancestral samples are picked at random and deemed to have a common parent. That memoryless property is the chief reason for its simplicity and usefulness.

However, the coalescent eliminates some important biological effects when the sample size is large relative to the population size. For example, if 10 haploid individuals are known to have one of two parents, the split of the two individuals between the parents is binomial in the Wright-Fisher (WF) model as we would expect. However, the partition of 10 individuals into two induced by the coalescent is uniform (Durrett, 2008) and not binomial.

The WF model, with its non-overlapping generations, is of course imperfect. For a phenomenon as complicated as the propagation of genetic material between generations and in diverse species, every model will be unsatisfactory in some way. The ability to make useful inferences by adjusting a few parameters is of greater consequence than microscopic faithfulness to actual phenomena, which are anyway unknowable to some extent. In this regard the WF model and even more so the Kingman model, with population sizes as well as mutation and recombination rates as their key parameters, have been quite useful.

The WF model is more reliable than the Kingman model when the sample sizes are large relative to population size. As implied by the classic birthday problem and its variants (Aldous, 1989), two individuals in a sample of size *N*^1/2^, assuming a fixed population size of 2*N*, may be expected to have a common parent. In samples of size *N*^1/2−*ϵ*^, *ϵ* > 0, there are no common parents in a typical generation in the limit of large *N*, and when there are common parents, it is reasonable to assume that at most two individuals have a common parent. The situation where every coalescence is a single double collision is the key assumption of Kingman’s coalescent. However, when the sample size is *N*^2/3^, three samples may be expected to have a common parent implying a triple collision. For sample sizes in-between *N*^1/2^ and *N*^2/3^, there will be simultaneous double collisions in a single generation. Kingman’s coalescent captures such simultaneous and multiple collisions as a succession of double collisions, each with no memory whatsoever of the previous collisions. Such an approximation introduces artifacts such as the uniform split of individuals between parents as we have already noted.

This paper derives theorems and algorithms that help delineate the regions of validity of Kingman’s coalescent relative to WF. Thus, the genealogical coalescence process is considered here using Kingman’s model as well as the WF model. The coalescent by itself will always refer to the Kingman model, although it will be explicitly referred to as the Kingman coalescent if there is room for ambiguity. Coalescence under the WF model is referred to as WF coalescence.

Existing literature on large sample effects may be divided into two categories. In the first category, the focus is on rates of coalescence and the number of ancestral samples as a function of the ancestral generation, and the Kingman model is assumed. Tavaré (1984) obtained formulas for the size of the ancestral sample (number of lineages) as a function of the ancestral generation, assuming fixed populations size. Griffiths and Tavaré (1998) obtained formulas that allowed the population size to vary. These formulas employ a sum whose terms alternate in sign and are inaccurate when the sample size is large, even assuming the coalescent approximation. Thus, Griffiths (2006) obtained asymptotic approximations that are better numerically for large samples. Other authors (Chen and Chen, 2013, Polanski et al., 2017) have extended this work to handle coalescence and inter-coalescence times. In particular, Chen et al. (2015) have observed that the number of segregating sites, an important statistic introduced by Watterson (1975) and which marked the shift from infinite alleles to the infinite sites model (Durrett, 2008), appears to be more robust under the coalescent approximation than the sample frequency spectrum for large sample sizes.

In the other category, the limitations of the coalescent are addressed explicitly using the WF model. Wakeley and Takahashi (2003) observed (relying on the earlier work of Fisher) that if the sample size is 2*Nx* with *x* ∈ (0, 1) (the same authors tweaked the WF model to allow even *x* ∈ (0, 2]), the expected parental sample size is 2*N*(1 − *e*^−*x*^). From that observation, they derived approximate estimates of the size of the coalescent tree as well as the length of the extremal branches. They concluded that large samples lead to more singletons (by about 10%) in the sample frequency spectrum. Fu (2006) used Stirling numbers to derive an exact coalescent for WF and came to a similar conclusion. More recently, Bhaskar et al. (2014) derived recurrences to compute the sample frequency spectrum as well as expectations of ancestral sample sizes exactly under WF. Connections to that work will be indicated later.

The contribution of this paper includes theorems that give sample sizes for which the coalescent agrees with WF (asymptotically). Simultaneous double collisions may be less disruptive (Bhaskar et al., 2014, Davies et al., 2007) than triple and other multiple collisions, and we prove theorems that indicate sample sizes for which all WF coalescences would be simultaneous double collisions and no worse. Algorithms that handle varying population sizes are derived and applied to demographic models of human population. These algorithms are used to visualize how bottlenecks in the distant past can disrupt the validity of the coalescent approximation. We prove a theorem about convergence of the sample frequency spectrum under WF to the classical answer derived using the coalescent. The Python/C program that implements algorithms we derive is posted at github.com/melfiand/lsample.

## Validity of the coalescent approximation for large samples

The coalescent consists of two independent stochastic processes (Kingman, 1982b). Let [*n*] denote the set {1,2,…, *n*}, which is the current sample. A partition of the set [*n*] is a set of nonempty subsets of [*n*] that are pairwise disjoint and whose union is the set [*n*]. In Kingman’s coalescent, the partition {*A*_1_,…, *A_k_*} of [*n*] is initialized to {{1},…,{*n*}} with *k* = *n*. At each step, two sets *A_i_* and *A_j_* are chosen, with each of the possible *k*(*k* − 1)/2 choices equally likely, and the two sets are replaced by their union *A_i_* ∪ *A_j_*. This stochastic process which governs the evolution of partitions of [*n*] is the first part of the coalescent and has been called the jump chain (Kingman, 1982b). A partition of [*n*] with *k* parts signifies an ancestral sample (in some earlier generation) of size *k*, with each ancestral sample denoted by the set of its descendants in the current sample. The merging of two partitions corresponds to two ancestral samples having a common parent so that the number of ancestral samples is reduced by 1.

The other part of the coalescent is the so-called death process (Kingman, 1982b), which governs the timing of the coalescence events. The death process is a continuous time Poisson process of varying rate, with the rate being *k*(*k* − 1)/2 when the number of ancestral samples is *k*. The connection to the WF model is made by equating a unit of time in the death process with 2*N* WF generations.

The jump chain and the death process are independent, and in many of our arguments, the death process does not play any role at all. The death process governs the rates of coalescence, which can be adjusted independently, for example to allow for varying population size, and at any rate the correspondence of the rates to the WF model is only approximate.

The following theorem of Kingman (1982b) characterizes the jump chain completely and does not depend upon the death chain:

*Suppose that the coalescent is run until the partition of* [*n*] *consists of exactly k sets. If the* |*A_j_*| = *n_j_ is the cardinality of A_j_*, *the probability that the partition into k sets is* {*A*_1_,…, *A_k_*} *is equal to*

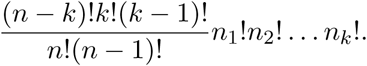

All italicized paragraphs are proved in the appendix. For the above statement, the appendix gives a combinatorial proof in the spirit of Griffiths and Lessard (2005). Kingman’s proof is recursive (Kingman, 1982b, Durrett, 2008).

The WF model says that if a haploid population of size *N*_1_ produces *N*_2_ children in the next generation, the split of the *N*_2_ children between *N*_1_ parents is multinomial (Durrett, 2008). When considering the genealogical process, the *k* samples in a generation choose parents from their parental generation independently, with each member of the parental generation being equally likely to be chosen. The members of the parental generation which turn out to be parents of any of the *k* samples constitute the parental sample. Such a passage from a sample to its parental sample will be referred to as a backward WF step. The WF genealogy of a sample is the sequence of backward WF steps until an ancestral generation with a single ancestral sample is reached.

The coalescent approximation may be violated because of simultaneous double collisions or triple collisions. Any multiple collisions higher than triple, such as quadruple collisions, always include triple collisions. Simultaneous double collisions in backward WF steps may be less disruptive because they can be produced by the coalescent with appreciable probability, as shown by the following corollary:

*Suppose the set*{{1},…,{*n*}} *undergoes k coalescences resulting in a partition into n* − *k sets. The probability q*(*k*, *n*) *that each set in the resulting partition is of size* 1 *or* 2 *is given by* 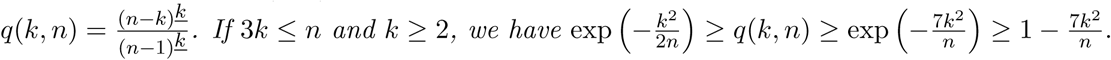

In this corollary, the falling power *n*(*n* − 1)… (*n* − *k* + 1) is denoted *n^<u>k</u>^* as recommended by Knuth (Graham et al., 1994, Knuth, 1997). The corollary implies that *k* simultaneous double collisions are produced with an appreciable probability as a result of *k* steps of the jump chain if *k* is much less than 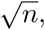 where *n* is the sample size. Therefore, we will look at bounds on *n* in terms of the population size 2*N* that allow only single double collisions in the WF genealogy of the sample (with high probability) as well as bounds that allow simultaneous double collisions.

For a constant population size of 2*N*, the following theorem indicates sample sizes that ensure that in every backward WF step at most two individuals have a common parent:

*The WF coalescent of a sample of size N*^1/3−*ϵ*^, *ϵ* > 0, *converges in probability to the coalescent in the limit of large N*.

This theorem does not consider rates of coalescence (encoded in the death process). The theorem only claims that the probability that there are either simultaneous double collisions or triple collisions in the WF genealogy of the sample goes to zero for large *N* for sample sizes smaller than *N*^1/3−*ϵ*^. However, for such sample sizes, the rates of WF coalescence agree with the rates of the coalescent (the death process) asymptotically, as will become clear from the statement and proof of a theorem about the sample frequency spectrum given below later.

Suppose we look for a bound on the sample size that ensures that every WF coalescence consists of either one or two double collisions. We then have the following theorem:

*The WF genealogy of a sample of size N*^2/5−*ϵ*^, *ϵ* > 0, *includes only one or two simultaneous double collisions in any ancestral generation with probability tending to* 1 *for large N*.

For another interpretation of this theorem, we may define the mod-2 coalescent in analogy with the Kingman coalescent. In an ancestral sample of size *k*, the mod-2 coalescent picks 4 individuals at random, divides them into two pairs, and merges both pairs. The merger can be thought of as a union of sets, with each set being the set of descendants present in the current sample of an individual in the ancestral sample. It is equivalent to ancestral individuals in both pairs finding common parents, the parents of the two pairs being distinct. The above theorem may then be interpreted as saying that the WF coalescent of samples of size *N*^2/5−*ϵ*^ or less is a mixture of the coalescent and the mod**-2** coalescent, with the proportion of the mixture varying with sample size.

More generally, we may allow *c* simultaneous double collisions rather than just 2. We have the following theorem:

*The probability that the WF genealogy of a sample of size* 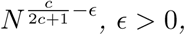 *consists only of double collisions*, *with the number of double collisions in any generation being one of*0, 1,…, *c*, *converges to 1 in the limit of large N*.

It is clear from this theorem that triple collisions kick in for sample sizes of the order *N*^1/2^ or higher. If *N* is large and the sample is size is smaller than *N*^1/2−*ϵ*^, we may assume that all collisions in backward WF steps are simultaneous double collisions.

Let *f* (*k*, *n*) be the probability that *k* out of *n* samples are mutants conditional on exactly one mutation in the genealogy of the sample. Let 𝓗_*n*_ denote the harmonic number 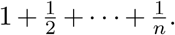 The coalescent implies 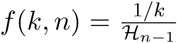 for *k* = 1,…, *n* − 1 in the limit of zero mutation rate. The following theorem shows that WF sample frequency spectrum converges to the classical answer for sample sizes smaller than *N*^1/3−*ϵ*^.

*Let fw F*(*k*,*n*) *be the probability that k out of n samples are mutants conditional on exactly one mutation in the WF genealogy of the sample. Then the total variation distance*

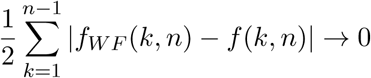

*for n* ≤ *N*^1/3−*ϵ*^, *ϵ* > 0, *in the limit of zero mutation and large N*.

## Algorithms for varying population sizes

For any sample size *n* > 2 and finite *N*, the probability that the WF genealogy of the sample includes simultaneous double collisions or triple collisions is strictly greater than zero. Indeed, the probability of such events in the transition to the parental generation by a single backward WF step is strictly greater than zero. However, by a theorem stated above and proved in the appendix, the probability that the WF genealogy includes only single double collisions converges to 1 in the limit *N* → ∞ if *n* ≤ *N*^1/3−*ϵ*^, where *N* is the constant population size.

The theorem may be amended to apply to populations sizes *CN*^1/3−*ϵ*^ for *ϵ* > 0 and some positive constant *C*. However, in the absence of a numerical value for the constant *C*, there would be no additional information.

A numerical constant can be produced by an algorithm that calculates the probability of the WF genealogy including only single double collisions. In this section, we derive such an algorithm. We derive another algorithm that calculates the probability that the genealogy of a sample of size *n* does not include any triple collisions. Both algorithms allow variable population sizes and may also be used to verify some of the asymptotic results.

Let *p*(0, *n*, *N*) be the probability that a sample of size *n* does not undergo any collision in a single Wright-Fisher step. Then

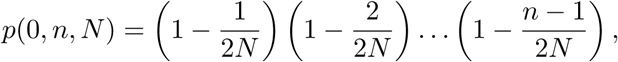

with 2*N* the population size of the parental generation. Let *p*(*k*, *n*, *N*) be the probability of exactly *k* double collisions and no triple collisions in a backward WF step with parental population size equal to 2*N*. Then

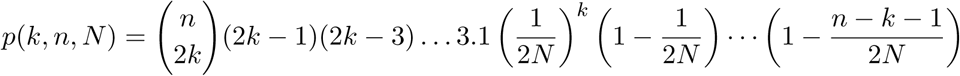

for 0 ≤ 2*k* ≤ *n*. The formula is valid for *n* > 2*N*. In fact, it is valid for any *k* with usual conventions about binomial coefficients (Graham et al., 1994) and with the assumption *n* ∈ ℤ^+^. The formula may be justified as follows. First, we may choose 2*k* samples to participate in *k* simultaneous double collisions in 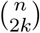 ways. To group the 2*k* samples into *k* pairs, the first of the chosen samples may be paired in (2*k* − 1) ways, the first of the remaining 2*k* − 2 samples may be paired in 2*k* − 3 ways, and so on. Thus, the total number of pairings is (2*k* − 1)(2*k* − 3)… 3.1. For each pair, the probability that the two samples in the pair have a common parent is 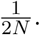 The remaining factors in the formula give the probability that the *k* pairs as well as the remaining *n* − 2*k* samples have *n* − *k* distinct parents.

### Probability of all collisions being single double collisions

For the current generation from which a sample of *n* is taken, we assume *t* = 0. Let 2*N*(*t*) be the haploid population size *t* ancestral generations ago. To calculate the probability that the WF genealogy of the sample has only single double collisions, the quantity *π*(*k*,*t*), *k* ∈ {1,…, *n*} is endowed with the following interpretation: at ancestral generation *t*, the probability that the ancestral sample is of size *k* with all prior coalescences being single double collisions is *π*(*k*, *t*). The allowed values for *k* are *k* = 1,…, 2*N*(*t*) for *t* > 0. When *k* = 0, however, *π*(*k*, *t*) has a different interpretation: *π*(0,*t*) is the probability that WF coalescence has produced something other than a single double collision prior to ancestral generation *t*. For *t* = 0, *k* can be anything, although the algorithm is initialized using *π*(*n*, 0) = 1 and *π*(*k*, 0) = 0 for *k* ≠ *n*, and in particular, *π*(*k*, 0) = 0 for 0 ≤ *k* ≤ *n* − 1.

Suppose the data at time *t* is *π*(*k*,*t*) with *k* ∈ [*n*] ∪ {0}. The crux of the algorithm is to generate data at time *t* + 1, and the recurrence

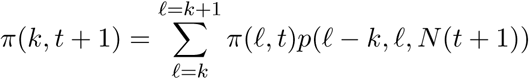

does that for *k* = 1,*…*, min(*n*, 2*N*(*t* + 1)). If the size of the ancestral sample in generation *t* + 1 is *k*, the ancestral sample size in generation *t* must be either *ℓ* = *k* or *ℓ* = *k* + 1 because simultaneous double collisions and triple collisions are precluded by the definition of *π*(*k*, *t* + 1). The two possibilities are disjoint, and the recurrence sums over the two possibilities.

The quantity *π*(0,*t* + 1), which has a different interpretation, is calculated using

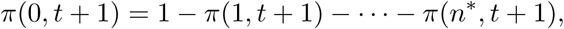

where *n*^∗^ = min(*n*, 2*N*(*t* + 1)).

The algorithm is terminated at the *t*th ancestral generation if *π*(0, *t*)+*π*(1, *t*) > 1 − 10^−4^. At termination, the probability that the sample has either coalesced to a single ancestral sample or the WF genealogy has violated the requirement of only single double collisions is greater than 0.9999.

### Probability of no triple collisions

The algorithm to calculate the probability of no triple collisions is similar. The quantity *π*(*k*, *t*), *k* ∈ ℤ^+^, now has the following interpretation: *π*(*k*,*t*) is the probability that the ancestral sample is of size *k* in ancestral generation *t* with no triple collision in any of the *t* backward WF steps from generation 0 to ancestral generation *t*. As before, the interpretation of *π*(0, *t*) is different: *π*(0, *t*) is the probability of a triple collision prior to ancestral generation *t*. Again as before, the algorithm is initialized using *π*(*n*, 0) = 1 and *π*(*k*, 0) = 0 for 0 ≤ *k* ≤ *n* − 1.

Suppose the data at time *t* is *π*(*k*, *t*) with *k* ∈ [*n*] ∪ {0}. The recurrence

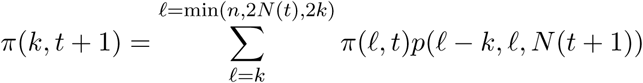

calculates data at *t* + 1 for *k* = 1,…, min(*n*, 2*N* (*t* + 1)). If the ancestral sample size at *t* + 1 is *k*, the ancestral sample size at *t*, which is denoted by *ℓ*, must be at least *k*. It can be at most 2*k* because any backward WF step that whittles down a sample of size greater than 2*k* to *k* must involve a triple collision. In addition, *ℓ* cannot exceed *n* or 2*N*(*t*). The recurrence is obtained by summing over all possibilities for *ℓ*. As before,

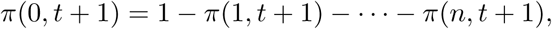

and we stop calculating when *π*(0, *t*) + *π*(1,*t*) > 1 − 10^−4^.

This algorithm can be sped up by ignoring *π*(*k*,*t*) if *π*(*k*,*t*) < *ϵ_tol_* for an *ϵ_tol_* that is small. As it is, the algorithm would maintain the probabilities *π*(*k*, *t*) for *k* ∈ [*n*] ∪ {0} typically. As *t* increases, a probability such as *π*(*n*,*t*) becomes quite small but can still remain positive. Holding on to such tiny numbers makes the algorithm quite expensive for large sample sizes. If probabilities smaller than *ϵ_tol_* are ignored, there is a rapid reduction in the sample sizes that are tracked at ancestral generation *t* in the initial stages of the algorithm if *n* is large. The total contribution of *π*(*ℓ*,*t*) to probabilities in all later stages is bounded by *π*(*ℓ*,*t*) because the recurrence sums over disjoint possibilities. Suppose all probabilities smaller than *ϵ_tol_* are ignored. The total probability ignored is then bounded by *ϵ_tol_nG*, where *n* is the sample size and *G* is the total number of generations. We use *ϵ_tol_* = 10^−120^ so that the ignored probability is vanishingly small even with *n* = *G* = 10^20^. A similar device was employed by Bhaskar et al. (2014).

## Verification and visualization

The algorithms for calculating the probability of WF coalescence with only single double collisions and no triple collisions enable a direct verification of the asymptotic theory. Figure 1 shows calculations for various population sizes. For each population size, the sample size at which the probability of coalescence according to the Kingman model (plot (a)) or with no triple collisions (plot (b)) are 5%, 50%, and 95% are also shown.

**Figure 1:**
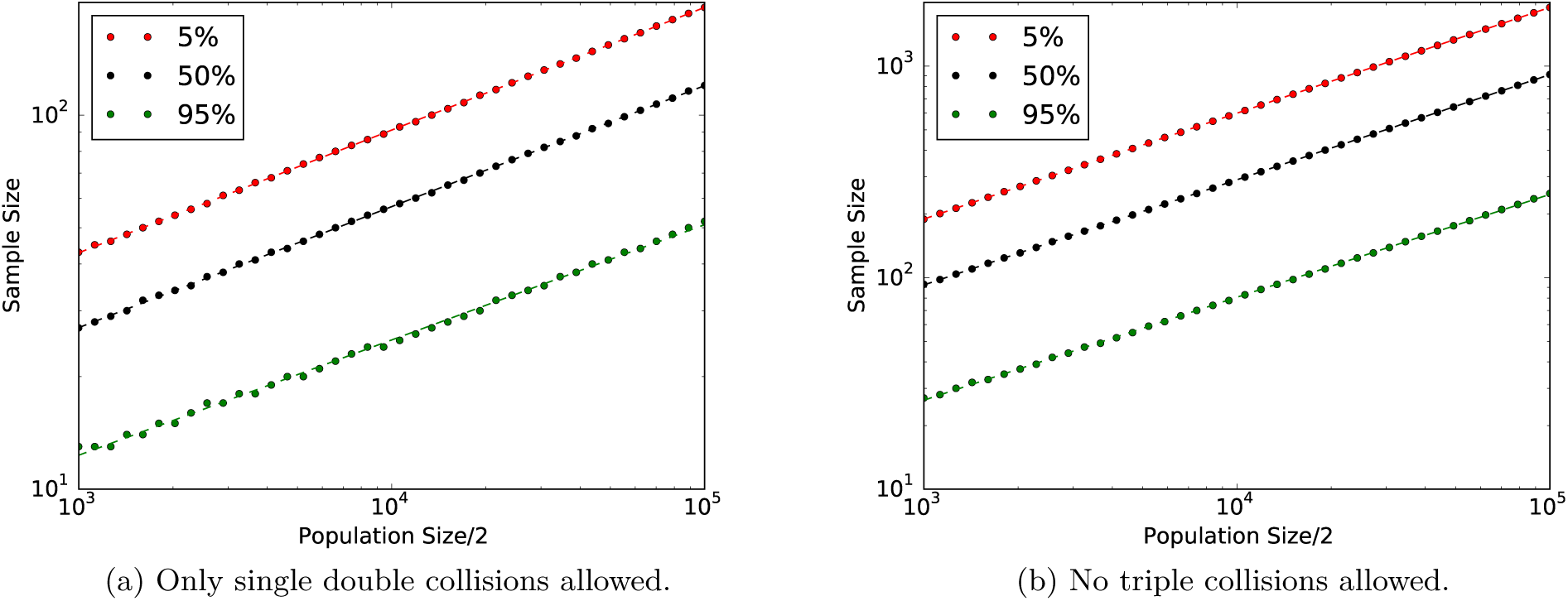
Exact probability of coalescence with only single double collisions and no triple collisions, respectively, for various constant population sizes. In each plot, the sample sizes at which the probability is 5%, 50%, and 95% are shown as solid circles. The dashed lines are linear fits.

Evidently, a higher sample size implies a higher probability of triple collisions or of violating the Kingman model. Sample sizes for which probabilities of conformance with the Kingman model are 5%, 50%, and 95% may be fitted as

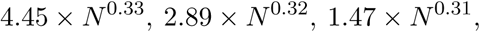

respectively. The quality of the fit is quite good for even *N* as small as 1000. The exponents are close to 1/3 as predicted by the asymptotic theory.

The quality fits for no triple collisions is even better. In this case, the sample sizes for which the probabilities of no triple collision are 5%, 50%, and 95% are

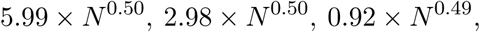

respectively. The exponents are close to 1/2 as predicted by the asymptotic theory. To increase the probability of WF coalescence with no triple collisions from 5% to 95% the sample size needs to be decreased by a factor of six approximately.

Both algorithms allow for variable population sizes. The four demographic models of human population we apply the algorithms to are the same as in Bhaskar et al. (2014). These models are:

- Constant population with *N* = 10^4^, which is the baseline assumption in human genetics (Durrett, 2008).
- Constant population with *N*(*t*) = 10^4^ except for two bottlenecks: the first being 620 < *t* ≤ 720 with *N*(*t*) = 500 and the second being 4620 < *t* ≤ 4720 with *N*(*t*) = 150, which is a drop-off by nearly a factor of 100. This model is based on Keinan et al. (2007).
- Exponential decay for 0 ≤ *t* ≤ 920 from *N*(0) = 3.5 × 10^4^ to *N*(920) = 10^3^, followed by *N*(*t*) = 2000 for 920 < *t* ≤ 2000, followed by *N*(*t*) = 15,000 for 2000 < *t* ≤ 5900, and *N*(*t*) = 6,500 for *t* > 5900. This model is based on Gravel et al. (2011). This model features a single exponential and is labeled `expl` in Figure 2.
- Exponential decay for 0 ≤ *t* ≤ 214 from *N*(0) = 5 × 10^5^ to *N*(214) = 10^4^, exponential decay for 214 ≤ *t* ≤ 920 with *N*(920) = 1025, *N*(*t*) = 2000 for 920 < *t* ≤ 2000, *N*(*t*) = 15000 for 2000 < *t* ≤ 5900, and *N*(*t*) = 6500 for *t* > 5900. This model features two exponentials and is therefore labeled `exp2` in Figure 2.

Figure 2 shows that the probabilities of triple collision and of violation of the Kingman model increase noticeably because of bottlenecks.

**Figure 2:**
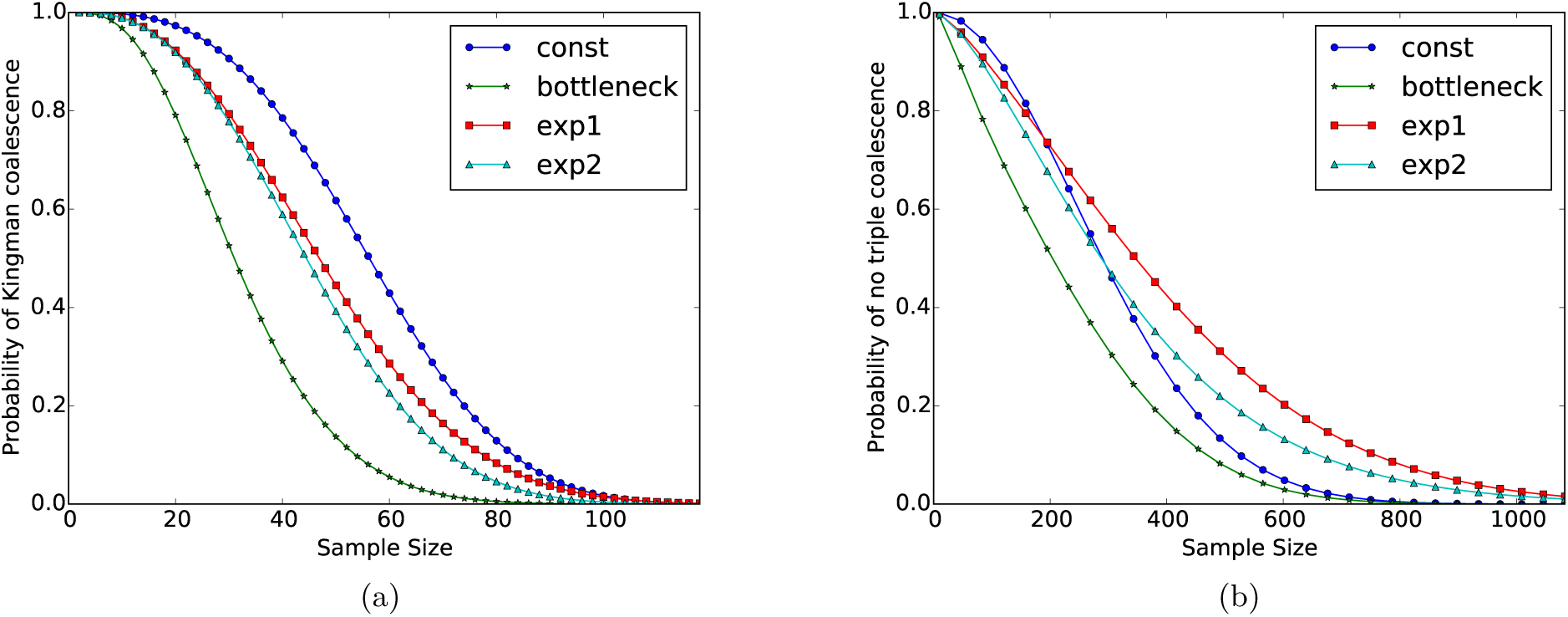
Probabilities of coalescence with at most a single double collisions in any backward WF step and with no triple collisions for four demographic models and various sample sizes.

Figures 3 and 4 give a more explicit visualization of the effect of bottlenecks. In Figure 3b, the distribution of possible ancestral sample sizes, conditioned on no violation, noticeably shifts downwards when the first bottleneck is encountered. The conditional probability of a violation spikes at the first bottleneck. At the second bottleneck, there is no such prominent spike in the conditional probability of a violation. However, the distribution of possible ancestral sample sizes, conditioned on no violation, noticeably shifts downwards at the second bottleneck, even though the bottleneck is more than 4500 ancestral generations away and the sample size is only 100.

**Figure 3:**
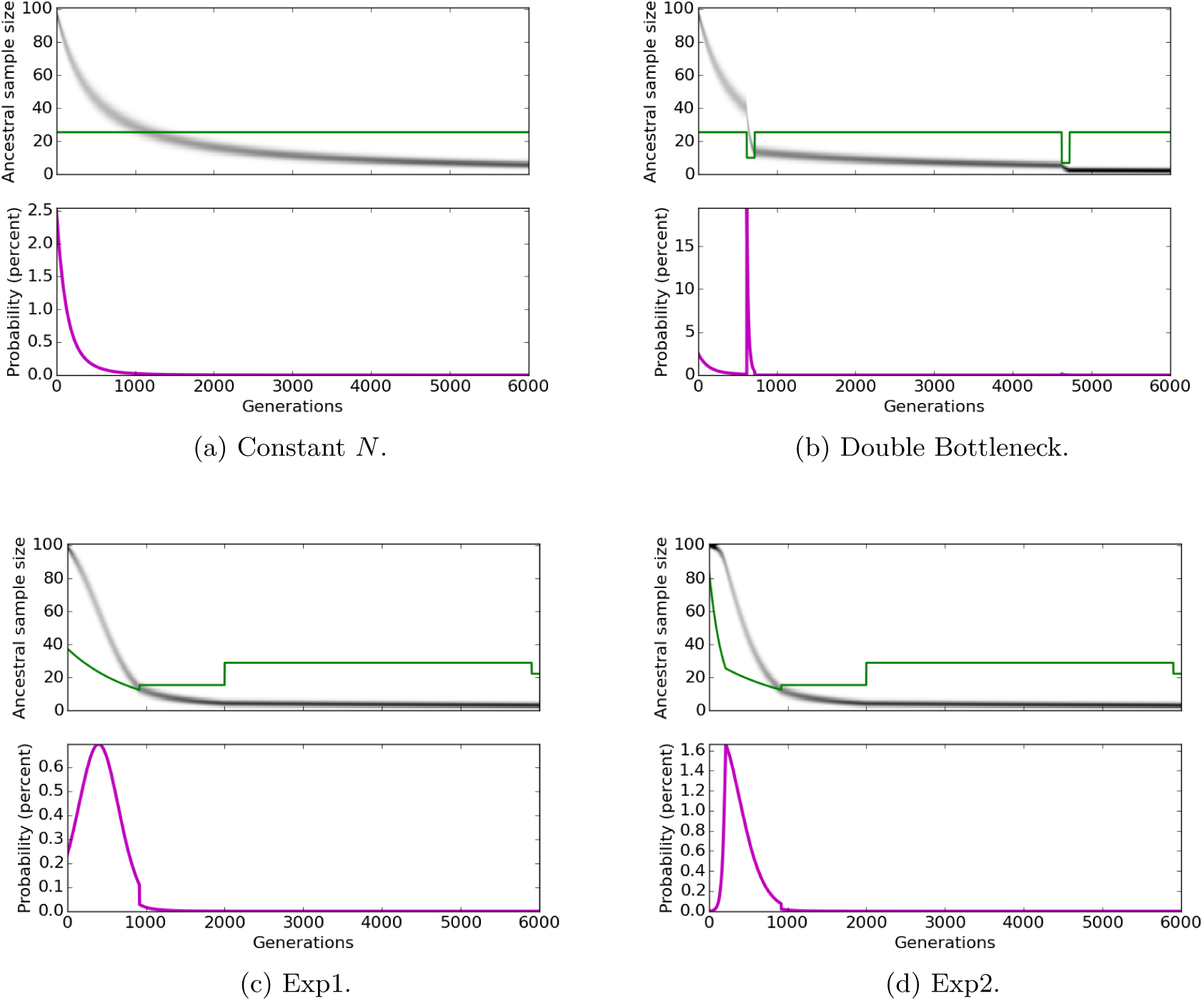
All four plots, which correspond to four different demographic models, are based on the algorithm to calculate the probability of coalescence allowing only single double collisions. In each case, the algorithm is applied with *n* = 100 at *t* = 0. Conditioned on no violation in any backward WF step from 0 to *t* (generations), there is a certain probability that the ancestral sample size at *t* is *k*. The upper panel is a heat-map of those probabilities, with black being 1 and white 0. The green line is a graph of 1.47 × *N*(*t*)^0.31^. The lower panel is a graph of the probability of a violation in the backward WF step from generation *t* to *t* +1 conditioned on no violation from 0 to *t*.

**Figure 4:**
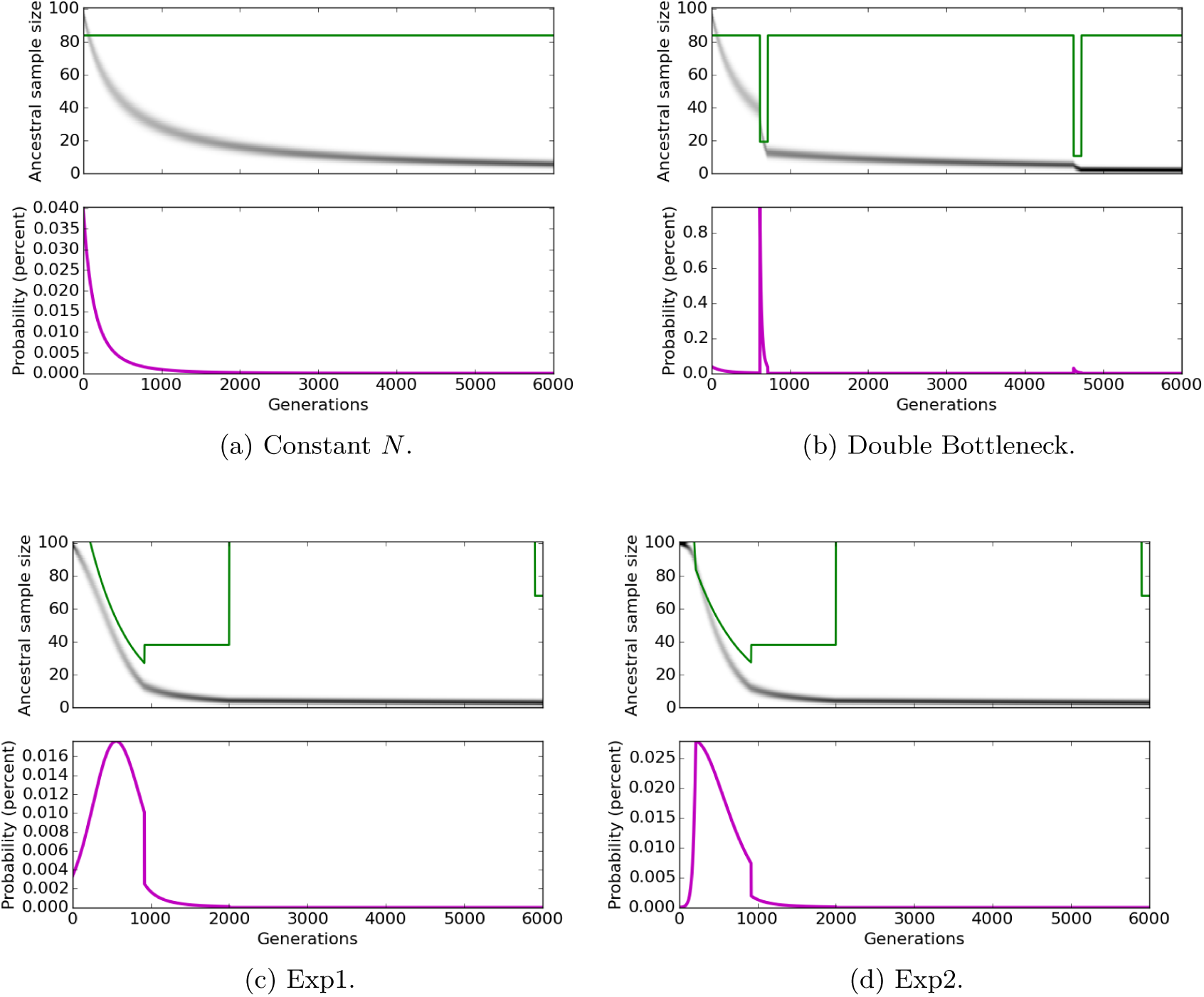
Same as Figure 3 except that the plots are based on the algorithm for calculating the probability of WF coalescence without triple collisions. The violations in this figure would be triple collisions, whereas the violations were simultaneous double collisions or triple collisions in the earlier figure. The green line in this figure graphs 0.92 × *N*(*t*)^0.49^.

Our interpretation of the phenomena in Figure 3 (c) and (d) is as follows. In both cases, the heat-maps of ancestral sample sizes (conditioned on no violation) show evidence of an inflection point. In these models with exponential decay in ancestral population sizes, there is less pressure on the sample to shrink initially. However, the exponential decay appears to eliminate that effect at the inflection point. In both plots, the spike in the conditional probability of a violation appears to be located near the inflection point.

In Figure 4, the same phenomena are in evidence, with a violation here being a triple collision and not a simultaneous double collision or a triple collision as in Figure 3. Some of the phenomena are a little more prominent here. For example, a small spike in the conditional probability of a violation is visible even at the second bottleneck in part (b) of the figure.

## Discussion

For a sample size of 60, the probability of deviation from the Kingman coalescent is higher than 50% in the human species. In the human species, the number of samples is often much greater than 60, and large sample effects cannot be ruled out.

What is the threshold beyond which large sample effects not captured by the Kingman coalescent are triggered? We have shown that the threshold is *N*^1/3^ in an asymptotic sense and that the asymptotic theory is fully relevant to realistic population sizes.

However, *N*^1/3^ is still a low threshold for the human species, and other studies (Bhaskar et al., 2014, Fu, 2006, Wakeley and Takahashi, 2003) have suggested that the sample frequency spectrum may not deviate too much for larger samples. We have shown that samples of size up to *N^1/2^* may experience simultaneous double collisions but not triple collisions in an asymptotic sense.

The sample frequency spectrum is only one of many possible statistics. Other statistics such as linkage disequilibrium may show deviations that are masked in the sample frequency spectrum. As noted by Kingman (1982b), the key to the coalescent is the probability distribution of the split of *n* samples between *k* ancestors in some past generation. Although this paper is developed from that point of view, the total variation distance between the probability distributions of the partitions under Kingman and WF is not yet known for large samples. Therefore, the question of how large the sample size must be to trigger effects that are not adequately handled by the Kingman model cannot be answered at the moment.

Nevertheless, there can be no doubt that the Kingman model produces unbiological effects for some large sample sizes, as we indicated in the introduction. Our computations show that even distant bottlenecks can impede the validity of the Kingman model. Non-African populations experienced bottlenecks during their first passage out of Africa and then again in later peregrinations around the globe. Whether unbiological effects of the Kingman model are to be accounted for in inferences of ancient bottlenecks is another interesting unanswered question.

## Appendix

This appendix gives proofs of theorems that were stated in the text. Statements of theorems are repeated in the interest of readability.

### Theorem 1

(Kingman (1982b)). *Suppose that the coalescent is run until the partition of* [*n*] *consists of exactly k sets. If the* |*A_j_*| = *n_j_ is the cardinality of A_j_*, *the probability that the partition into k sets is* {*A*_1_,…, *A_k_*} *is equal to*

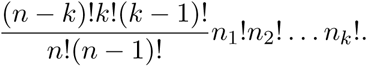

*Proof.* Because each coalescence is a union of two disjoint subsets of [*n*], the coalescent process can be depicted as a forest of binary trees with each vertex a subset of [*n*] and with the leaves being {1},… {*n*}. If disjoint subsets *S*_1_ and *S*_2_ coalesce, then *S*_1_ ∪ *S*_2_ occurs as a vertex with *S*_1_ and *S*_2_ as its two children. Coalescences deeper into the ancestry are placed higher to capture the ordering of events. The leaves are lowest, and no two interior vertices occur at the same height. Because the Kingman coalescent is memoryless, every coalescent tree with the same root is generated with the same probability.

The number of coalescent trees with root *A*_1_ and with their *n*_1_ leaves being equal to {*j*} for *j* ∈ *A*_1_ is equal to 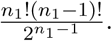 That is because the first union is any one of *n*_1_(*n*_1_ − 1)/2 possibilities, the second union any one of (*n*_1_ − 1)(*n*_1_ − 2)/2 possibilities, and so on. The total number of coalescence events in any of these trees is *n*_1_ − 1. Likewise, the number of coalescent trees with root *A_j_* is 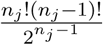 and the number of coalescence events in any of these trees is *n_j_* − 1.

The total number of forests with roots equal to *A*_1_,…,*A_k_* is equal to

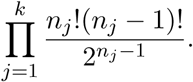

Although the order of coalescence events within a single tree is determined, the order of events between different trees is not determined. Because the number of coalescence events in a tree with root *A_j_* is *n_j_* − 1, the coalescence events corresponding to any given forest can be ordered in

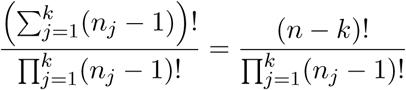

ways. Thus, the total number of sequences of *n* − *k* coalescence events resulting in a forest with roots *A*_1_,…,*A_k_* is equal to 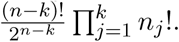 Each sequence of (*n* − *k*) coalescence events is equally likely by the memoryless property of the coalescent, and the total number of sequences of length *n* − *k* is equal to 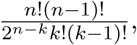 which implies the stated theorem.

### Corollary 2.

*Suppose the set* {{1},…, {*n*}} *undergoes k coalescences resulting in a partition into n* − *k sets. The probability q*(*k*, *n*) *that each set in the resulting partition is of size* 1 *or* 2 *is given by* 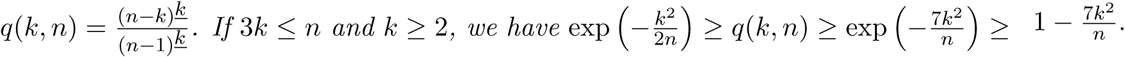

*Proof.* The probability *q*(*k*, *n*) is zero if 2*k* ≥ *n* + 1 because a partition of size 3 or more is inevitable after so many coalescences. The formula for *q*(*k*, *n*) is easily verified in this case.

Now suppose 2*k* ≤ *n*. If a partition into *n* − *k* sets has only sets of sizes 1 and 2, the number of sets of sizes 1 and 2 must be (*n* − 2*k*) and *k*, respectively. The number of such partitions is equal to

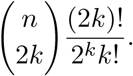

By Theorem 1, the probability of each partition is equal to

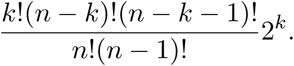

The proof of the formula for *q*(*k*,*n*) is completed by multiplying the two numbers and simplifying.

The stated bounds for *q*(*k*, *n*) follow from calculations that are elementary but a little tedious.

Let *p* = *q*(*k*,*n*). To bound *p*, note that 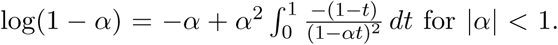 If 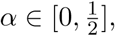 we may deduce that

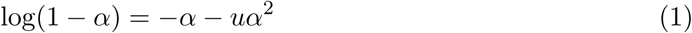

for some 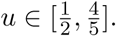 By similarly elementary arguments,

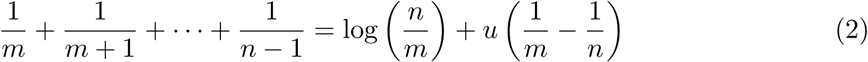

and

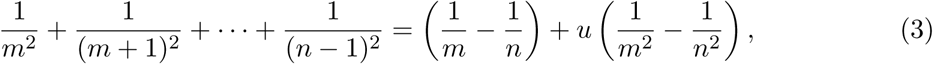

for *m*,*n* ∈ ℤ^+^, *m* < *n*, and some *u* ∈ [0, 1].

From the formula for *p* and (1), we have

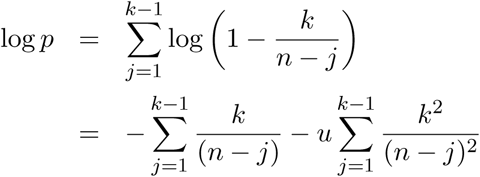

for some 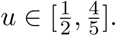 The application of (1) is justified because 3*k* ≤ *n* implies *k/*(*n*−*k*+1) < 1/2. Applying (2) and (3), we get

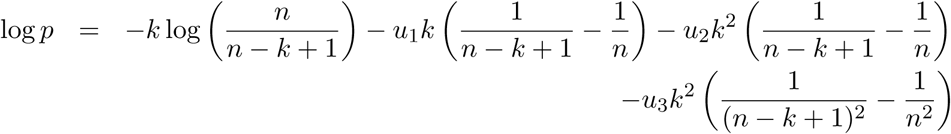

for some 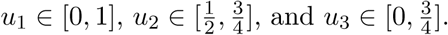

Thus,

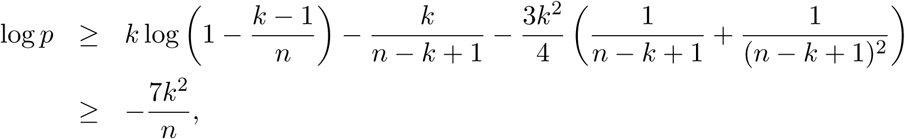

where the second inequality is obtained using (1). We then have *p* ≥ exp(−7*k*^2^/*n*) ≥ 1 − 7*k*^2^/*n*, proving the lower bound.

To prove the upper bound, argue as follows:

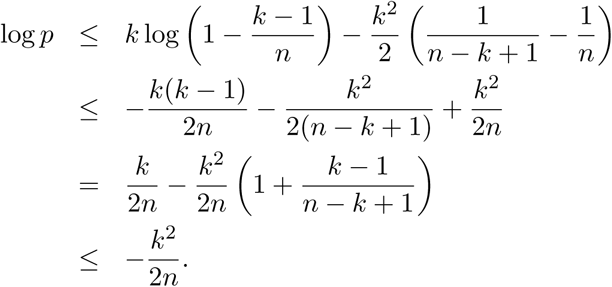

### Lemma 3.

*Consider the application of a single backward WF step to a sample of size n with parental population of size 2N. Let p_d_ be the conditional probability that there is a single double collision given that there is a collision. Then*

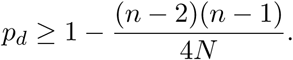

*Proof.* Let *A*_12_ be the event that samples 1 and 2 collide (in one generation), more specifically 1 and 2 have the same Wright-Fisher parent. Obviously 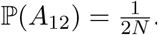.

Let 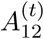 be the event that 1 and 2 collide and that one of the other (*n* − 2) samples has the same parent as 1 and 2, implying a triple collision or worse. We have 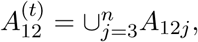 where *A*_12*j*_ is the event where 1, 2, and *j* have the same parent. Because 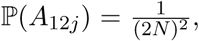 we have 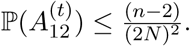

Let 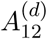 be the event that 1 and 2 collide and that there is some other pair that collides, implying two double collisions or worse. We have 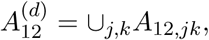 where *A*_12,*jk*_ is the event that 1,2 as well as *j*,*k* have the same parent. The union is over 2 < *j* < *k* − *n*. Because 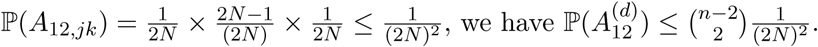

If *Ã*_12_ is the event that 1,2 collide and there are no other collision, 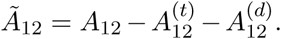 Therefore,

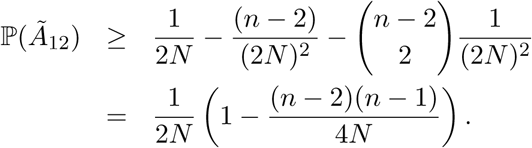

The probability that there is a single double collision during a backward Wright-Fisher step is equal to 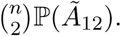

If 𝓒 is the event that there is some collision, 𝓒 = ∪_*j*,*k*_*A_jk_*, union over 1 ≤ *j* < *k* ≤ *n*. Therefore, the probability of a collision 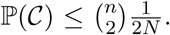 The lower bound for *p_d_* in the lemma is obtained by simplifying (2*N*)ℙ(*Ã*_12_).

### Theorem 4.

*The WF genealogy of a sample of size N*^1/3−*ϵ*^, ϵ > 0, *includes at most one double collision in any given generation*, *with probability converging to* 1 *as N* → ∞.

*Proof.* Let 𝓓 be the event that a sample of size *n* undergoes more than a single double collision in some backward WF step before coalescing to a single ancestor. Let 𝓓_*k*_ be the event that the ancestral sample size is equal to *k* in some generation but the ancestral sample size is never *k* − 1. Evidently, 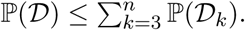

By Lemma 3,

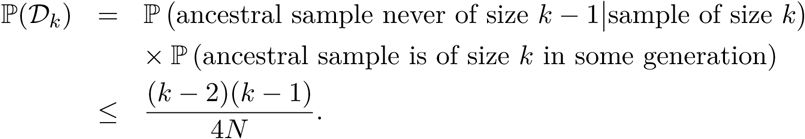

Therefore,

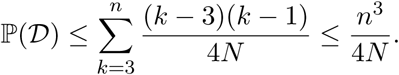

If *n* = *N*^1/3−*ϵ*^, ℙ(𝓓) ≤ *N*^−3*ϵ*^/4, which converges to zero as *N* → ∞. In the complement of 𝓓, WF coalescence follows the Kingman model proving the theorem.

Let 𝓓_*n*_ be the event that there are more than two double collisions or a triple collision or worse. Then

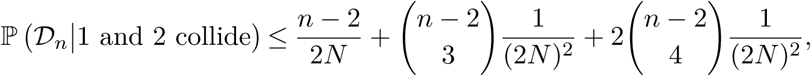

where the first term accounts for any of the samples 3 through *n* having the same parent as 1 and 2, the second term accounts for triple collisions with a parent other than that of 1 or 2, and the third term accounts for the possibility that there are two or more additional double collisions. This bound simplifies to

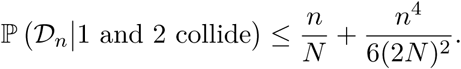

This is an almost correct bound for the conditional probability of 𝓓_*n*_ given *any* collision, as we may expect from the high degree of symmetry. The argument below makes the idea rigorous by using more detailed conditioning.

The event 𝓒_*j*,*k*_, which we presently define and with respect to which we will condition later, pertains to a single backward WF step. The sample size is assumed to be *n*.

- Samples 1 through *j* − 1 have unique parents and do not collide with any sample.
- The parent of sample *j* differs from the parents of samples *j* + 1,…, *k* − 1.
- Samples *j* and *k* have the same parent.

### Lemma 5.

*Let p*_1_ *be the conditional probability given 𝓒_j_*,*k that some sample has the same parent as j and k. We have*

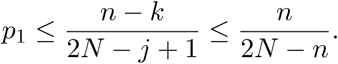

*Proof.* None of the samples [*k*] − {*j*, *k*} are allowed to have the same parent as *j* and *k* subject to the condition 𝓒_*j*,*k*_. Subject to the condition 𝓒_*j*,*k*_, samples *k* + 1,…,*n* can have any of 2*N* − *j* + 1 parents (the *j* − 1 parents of 1,…,*j* − 1 are excluded). The probability of ending up with the same parent as *j* and *k* is thus 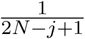 for each of those *n* − *k* samples, which proves the lemma

### Lemma 6.

*Let p*_2_ *be the conditional probability given* 𝓒_*j*,*k*_ *that some three samples have the same parent and that that parent is distinct from the parent of j and k. We have*

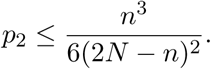

*Proof.* The three samples of this lemma cannot belong to [*j*] ∪ {*k*}. Thus, the three samples must be chosen out of a set of cardinality *n* − (*j* + 1), which can be done in

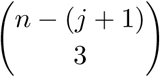

ways. For any such choice, the probability of a triple collision given 𝓒_*j*,*k*_ is

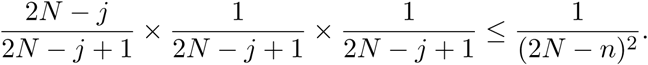

The first factor accounts for the first member of the triple having to choose a parent other than those of [*j*] and 1/(2*N* − *j* + 1) is the probability that the second or third member choose the same parent as the first, subject to 𝓒_*j*,*k*_. Therefore,

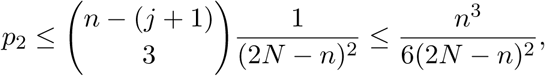

as claimed in the lemma.

### Lemma 7.

*Let p*_3_ *be the conditional probability given* 𝓒_*j*,*k*_ *that for each of c or more pairs*, *the two members of the pair have a common parent with that parent being distinct from the parents of all other pairs as well as the parent of j and k. We have*

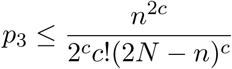

*Proof.* The *c* pairs must be chosen out of the samples [*n*] − [*j*] − {*k*}. That means *n* − (*j* + 1) choices for each member of a pair and the samples which form *c* pairs can be chosen in

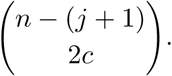

Having chosen the 2*c* samples, they can be paired in

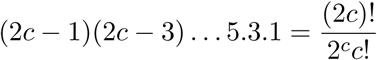

ways because the first of the chosen samples can be paired in 2*c* − 1 ways following which the second of the remaining samples can be paired in 2*c* − 3 ways and so on. Having formed the pairs, the probability given *C*_*j*,*k*_ that each pair has a common parent distinct from that of other pairs as well as *j* and *k* is

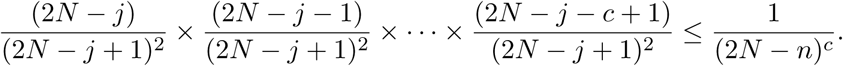

Therefore,

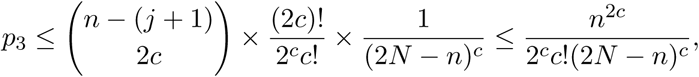

as claimed in the lemma.

### Lemma 8.

*Let* 𝓓_*n*_ *denote the event that there are c* + 1 *or more double collisions with distinct parents or some triple collision in a single backward WF step applied to a sample size of n. Then*

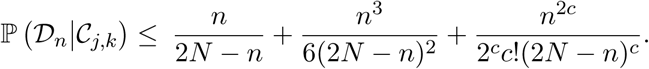

*Proof.* The event 𝓓_*n*_ ∩ *C*_*j*,*k*_ implies one of the following:

- Some sample has the same parent as *j* and *k*.
- Some three samples have a common parent distinct from the parent of *j* and *k*.
- There are *c* or more double collisions in addition to the collision between *j* and *k*.

Therefore, ℙ(𝓓_*n*_|*C*_*j*,*k*_) ≤ *p*1 + *p*_2_ + *p*_3_ proving the lemma.

### Theorem 9.

*The WF genealogy of a sample of size* 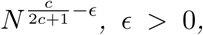 *includes at most c simultaneous double collisions and no triple collisions with probability converging to* 1 *in the limit of large N*.

*Proof.* Let 𝓓_*ℓ*_ be the event that the ancestral sample size is *ℓ* and a backward WF step results in either a triple collision or more than *c* double collisions. From the previous lemma,

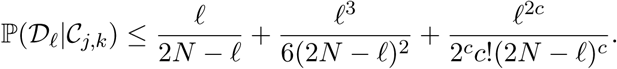

Let 𝓒 denote the event that a collision has occurred in a backward WF step with a sample size of *ℓ*. Evidently, 𝓒 is the disjoint union of the events 𝓒_*j*,*k*_ over 1 − *j* ≤ *k* − *ℓ*, with the event 𝓒_*j*,*k*_ asserting the first collision in lexicographic order is between sample *j* and *k.* Therefore,

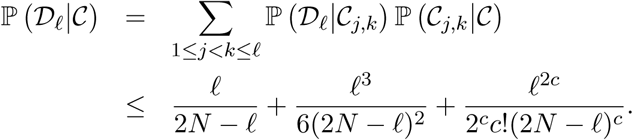

Let 𝓓 be the event that a sample of size *n* undergoes either a triple collision or more than *c* double collisions in some generation before coalescing to a single ancestor under WF. Then

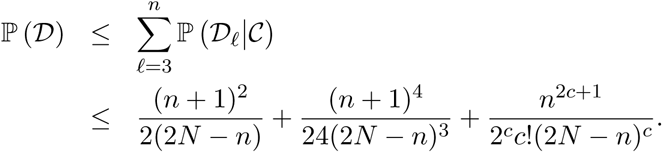

The proof is completed by substituting 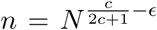 and verifying the *N* → ∞ limit to be zero.

We now turn to the sample frequency spectrum under WF. Let *q*(*n*, 2*N*) denote the probability of a coalescence event in a single backward WF step. Then

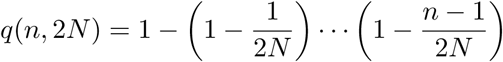

The probability of a mutation event in a single backward WF step is assumed to be *nμ*. Given that either a mutation event or a coalescence event has occurred, the probability that it is a mutation is equal to

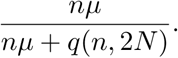

The probability it is a coalescence is equal to

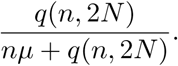

We are making the usual assumption that the sample cannot be hit with both a mutation and a coalescence in the same generation. The assumption would be unreasonable for large samples. However, we limit ourselves to samples of size *N*^1/3−*ϵ*^ or less. In addition, we condition to limit the total number of mutations in the genealogy of the sample to one, which makes the assumption reasonable even for large *N*.

In particular, denote the event that the genealogy of a sample of size *n* (in either WF or Kingman model) includes exactly one mutation by 𝓒. The probability that the mutation strikes when the WF ancestral sample size is *k* is equal to

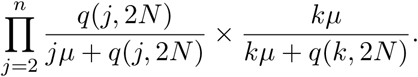

Therefore, conditioned on 𝓒 the probability that mutation strikes a sample of size *n* before any coalescence event is equal to

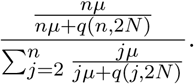

We take the limit *μ* → 0 to get

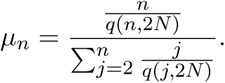

Thus, *μ_n_* is the probability that a mutation is the first event to strike a sample of size *n* conditioned on 𝓒.

Let *f* (*j*,*n*) be the probability that *j* out of *n* samples are mutants conditioned on 𝓒. The recurrence for *f* (*j*, *n*) is

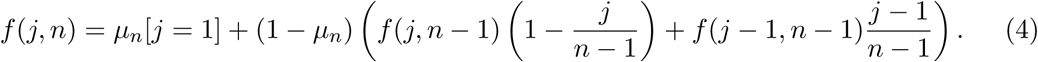

In this recurrence, we have used Knuth’s notation (Graham et al., 1994, Knuth, 1997) by which [*j* = 1] evaluates to 1 if *j* = 1 and 0 otherwise. To obtain the classical formula for the sample frequency spectrum, replace *μ_n_* by

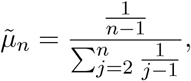

which is obtained by taking *q*(*j*, 2*N*) = *j* (*j* − 1)/4*N* following the Kingman model. The exact solution of the recurrence

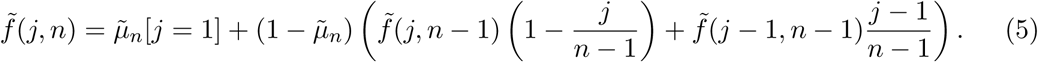

is given by

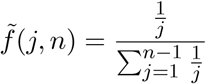

for *j* = 1,…, *n* − 1.

### Lemma 10.

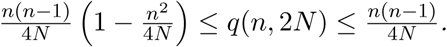

*Proof.* The proof of Lemma 3 shows that 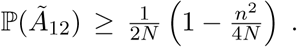 The proof of the lower bound follows from 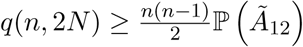 and the upper bound is obviously true.

### Lemma 11.

*For n*^2^ ≥ 2*N*,

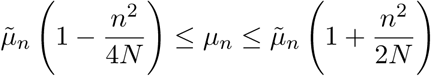

*for n*^2^ ≤ 2*N*.

*Proof.* If we use the definition of *μ_n_* and write

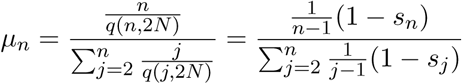

after taking 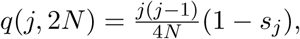 then by the previous lemma *S_j_* ∈ [0,*j*^2^/4*N*].

To obtain the lower bound in the lemma, set *S_j_* = 0 in the denominator and *s_n_* = *n*^2^/4*N* in the numerator.

To obtain the upper bound in the lemma, set *S_n_* = 0 in the numerator and replace *S_j_* by 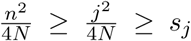 in the denominator. Finally, observe that 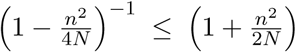 if *n*^2^ ≤ 2*N*.

### Lemma 12.

*If n* ≤ *N*^1/3−*ϵ*^, *then*

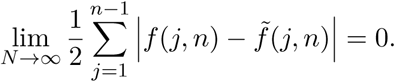

*Proof.* Note that |*ab* − *ãb̃*| ≤ |*a* − *ã*| |*b*| + |*b*| − *b̃*| |*a*|. Subtracting (4) and (5), we get

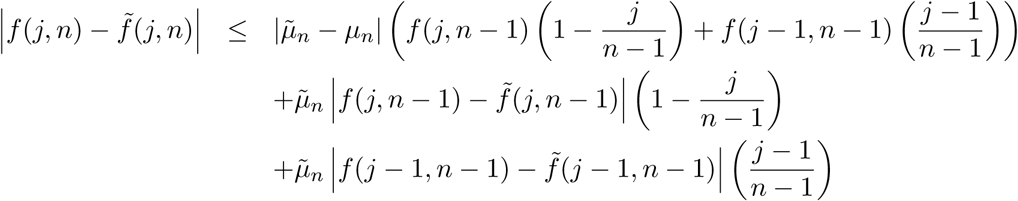

for *j* = 2,…,*n* − 1. For *j* = 1, there is an additional |*μ_n_* − *μ̃_n_*| term.

Summing these inequalities, we have

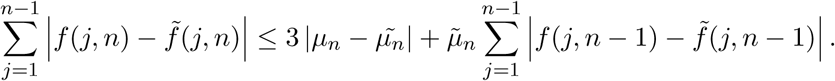

Because *μ̃_n_* < 1 for *n* > 2 and *f* (*j*, *n*) = *f̃*(*j*, *n*) for *n* = 2, we have

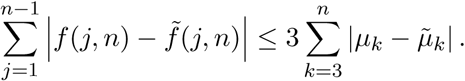

The proof is now easily completed by an application of the previous lemma.

### Theorem 13.

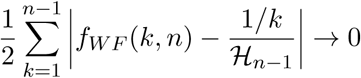

*for n* ≤ *N*^1/3−*ϵ*^, *ϵ* > 0, *in the limit of zero mutation and large N*.

*Proof.* By Theorem 4, the probability that any backward WF step produces a simultaneous double collision or a triple collision converges to zero as *N* → ∞. Thus, we may invoke the previous lemma and infer this theorem.

## Footnotes

1 The “ancestral sample” nomenclature is more intuitive for our purposes. However, in the context of the coalescent, the same concept is referred to as “lineage” or “ancestral lineage” (Griffiths, 2006, Griffiths and Tavaré, 1998, Tavaré, 1984).

## References

D. Aldous. Probability Approximations via the Poisson Clumping Heuristic. Springer, New York, 1989.

A. Bhaskar, A. G. Clark, and Y. S. Song. Distortion of genealogical properties when the sample is very large. Proceedings of the National Academy of Sciences, 111:2385–2390, 2014.

H. Chen and K. Chen. Asymptotic distributions of coalescence times and ancestral lineage numbers for populations with temporally varying size. Genetics, 194:721–736, 2013.

H. Chen, J. Hey, and K. Chen. Inferring very recent population growth rate from population-scale sequencing data: using a large-sample coalescent estimator. Molecular Biology and Evolution, 32:2996–3011, 2015.

J. L. Davies, F. Simančík, R. Lyngsø, T. Mailund, and J. Hein. On recombination-induced multiple and simultaneous coalescent events. Genetics, 177:2151–2160, 2007.

R. Durrett. Probability models for DNA sequence evolution. Springer Science & Business Media, 2008.

W. J. Ewens. The sampling theory of selectively neutral alleles. Theoretical Population Biology, 3:87–112, 1972.

Y. Fu. Exact coalescent for the Wright-Fisher model. Theoretical Population Biology, 69: 385–394, 2006.

R.L. Graham, D.E. Knuth, and O. Patashnik. Concrete Mathematics. Addison-Wesley, NJ, 2nd edition, 1994.

S. Gravel, B. M. Henn, R. N. Gutenkunst, A. R. Indap, G. T. Marth, A. G. Clark, F. Yu, R. A. Gibbs, C. D. Bustamante, D. L. Altshuler, et al. Demographic history and rare allele sharing among human populations. Proceedings of the National Academy of Sciences, 108: 11983–11988, 2011.

R.C. Griffiths. Coalescent lineage distributions. Advances in Applied Probability, 38:405–429, 2006.

R.C. Griffiths and S. Lessard. Ewens’ sampling formula and related formulae: combinatorial proofs, extensions to variable population size and applications to ages of alleles. Theoretical Population Biology, 68:167–177, 2005.

R.C. Griffiths and S. Tavaré. The age of a mutation in a general coalescent tree. Stoch. Models, 14:273–295, 1998.

A. Keinan, J. C. Mullikin, N. Patterson, and D. Reich. Measurement of the human allele frequency spectrum demostrates greater genetic drift in east asians than in europeans. Nature Genetics, 39:1251, 2007.

J. F. C. Kingman. On the genealogy of large populations. Journal of Applied Probability, 19: 27–43, 1982a.

J. F. C. Kingman. The coalescent. Stochastic Processes and their Applications, 13:235–248, 1982b.

D.E. Knuth. The Art of Computer Programming, volume 1. Addison-Wesley, NJ, 3nd edition, 1997.

A. Polanski, A. Szczesna, M. Garbulowski, and M. Kimmel. Coalescence computations for large samples drawn from populations of time-varying sizes. PloS One, 12, 2017.

S. Tavaré. Line-of-descent and genealogical processes, and their applications in population genetics models. Theoretical Population Biology, 26:119–164, 1984.

J. Wakeley and T. Takahashi. Gene genealogies when the sample size exceeds the effective size of the population. Molecular Biology and Evolution, 20:208–213, 2003.

G. A. Watterson. On the number of segregating sites in genetical models without recombination. Theoretical Population Biology, 7:256–276, 1975.

